# Biomechanical response of the human brain to low-intensity blast: a finite element study of single and repeated exposures

**DOI:** 10.64898/2026.07.25.740699

**Authors:** Melissa Dunphy Yates, Thomas A. Metzger, Vanessa D. Alphonse, Kyle A. Ott, Eyal Bar-Kochba

## Abstract

Repetitive low-intensity blast (LIB) exposure has been identified as a probable cause of mild blast-induced traumatic brain injury (mbTBI) and a chronic injury risk to U.S. military personnel. However, the human brain’s biomechanical response to this loading regime remains poorly characterized. Using blast-exposure-validated 3D models of human anatomy, we simulated the intracranial tissue response to blast pressure typically experienced by Warfighters during weapons training. Two scenarios were evaluated, a single-dose exposure and a repetitive-dose exposure, to study intracranial pressure (ICP), shear strains, and spectral content. Ansys LS-DYNA was used to generate planar blast waves with peak overpressures of 4–90 kPa and positive phase durations of 2.2–10 ms. Single exposures produced ICP ranging from 4.7–112.7 kPa, dependent on dose and positive phase duration. Under repetitive LIB exposure, peak ICP increased by 8–26% relative to single exposures, with an increase of high-frequency components (*>*2 kHz). These results demonstrate that LIB can produce measurable intracranial responses that are amplified through repetition, producing pronounced spectral content and elevated pressures despite low strain levels. This study underscores the need to further investigate cumulative dose effects and the value of computational approaches to clarify hypothesized mbTBI mechanisms in operationally relevant conditions.

## Introduction

Military and law enforcement personnel are routinely exposed to repetitive low-intensity blast (LIB) during training and combat while operating weapon systems, including artillery, mortars, shoulder-fired arms, stun grenades, and breaching explosives [1–6]. Dosimetry studies have reported blast overpressures during these events of up to 90 kPa, substantially exceeding the DoD’s interim blast-overpressure safety threshold of 28 kPa (4 psi) for military personnel [7]. During a single training exercise, personnel can be exposed to more than 100 LIB events, [8] leading to numerous exposures throughout their careers. While extensive research has focused on blast-induced traumatic brain injury (bTBI) caused by high-level blast explosions, [9] emerging evidence indicates that LIB exposure can lead to subconcussive or mild blast-induced traumatic brain injury (mbTBI), [4, 10] which is associated with cognitive deficits, anxiety-related disorders, sleep disturbances, reduced sociability, and impaired motor function [4, 10–13]. Blast exposure and mTBI have been associated with persistent neurological changes and an increased risk of long-term neurodegenerative outcomes [14–16].

Despite more than a decade of research, the injury mechanisms for mbTBI are still poorly understood, particularly for repetitive LIB. The pathophysiology of mbTBI has highlighted structural (e.g., diffuse axonal injury) and biochemical abnormalities in the brain, spinal cord, and nerves of mbTBI patients [2, 17–21]. However, linking brain biomechanics to these pathophysiological effects is essential to clarify mbTBI mechanisms as a result of these complex and repetitive loads.

Animal, cadaveric, and synthetic physical surrogate models have been used to study brain biomechanics under blast. While these efforts have provided insights into the biomechanics of mbTBI, these models each have limitations for direct translation to military and law enforcement personnel. Animal models link intracranial responses to pathophysiology, but anatomical differences lead to a mismatch of boundary conditions, limiting translation to humans [22–24]. Cadaver models resolve anatomical issues but lack living pathophysiology and suffer from brain material degradation [25]. Finally, synthetic physical surrogate models face anatomical, material, and pathophysiological limitations, but provide ease of testing in blast environments [26–28]. Taken together, experimental models alone do not sufficiently capture the biomechanical response of the brain to blast exposure.

Computational models offer unique insights into the biomechanical response of the brain to blast-overpressure exposures. Through recreation of the physics associated with blast overpressure and incorporation of the complex human anatomy, geometry, and material properties, these models are able to quantify the spatiotemporal pressure-wave transmission in the brain, typically unachievable by experimental models. Previous work has predicted that peak intracranial pressure (ICP) is approximately 30% less than the reflected pressure at the surface of the blast impact, with brain strains less than 10% and highly dependent on the impulse of the blast [25, 29]. Additionally, Tan et al. investigated Warfighter occupational blast exposures; specifically, characterizing the role of head posture positioning and repetitive exposures on cumulative brain response [30]. This study in particular is one of the first computational efforts to characterize the cumulative effect of repetitive brain loading due to Warfighter occupational exposures. While these studies have provided significant insights into the blast intracranial biomechanical environment, they do not sufficiently encompass the range of occupational blast exposures measured during recent training exercises [8]. Indeed, characterization of the intracranial biomechanical response to repetitive, low-magnitude occupational blast exposures remains a gap in understanding mbTBI injury mechanisms and risks.

In seeking to address this gap, the current study presents a numerical model-based approach using a validated human head-brain finite element (FE) computational model leveraging recently collected blast overpressure data during military training exercises [8] to quantify the brain biomechanical response to both single and repetitive LIB exposures. Given the previously observed geometric and material attenuation of blast within the brain, we hypothesize that repetitive LIB exposure will result in altered wave transmission and increased localized ICP. The results of this work aim to elucidate the potential biomechanical injury mechanisms associated with mbTBI and can help support future *in vitro* and *in vivo* studies linking LIB to the pathophysiology.

## Methods

### Human head modeling

Ansys LS-DYNA R14.0.0 (Ansys, Inc., Canonsburg, PA, USA), a commercial finite element analysis software, was employed to computationally model the human biomechanical response to LIB exposure. A fluid-structure coupling approach using the arbitrary Lagrangian–Eulerian (ALE) formulation was developed. The air domain and detonation were modeled as Eulerian elements and the three-dimensional (3D) human head anatomy was modeled as FE Lagrangian solids to capture BOP transmission within the brain. This coupled approach enabled accurate pressure transmission from the BOP wave to the human anatomy. The Applied Physics Laboratory (APL) Head-Neck model used here was previously described. [31] The model mesh was created in TrueGrid (XYZ Scientific Applications, Inc., Pleasant Hill, CA) to reflect the 50th percentile male based on the ANSUR II and Visible Human geometries. [32, 33]

The APL Head-Neck model included anatomically detailed representations of the brain, cerebrospinal fluid, dura mater, falx cerebri and tentorium cerebelli, cortical and trabecular cranial bone, sinuses, flesh, skin, bridging veins, and cervical spine structures (Fig. 1). Component-specific element counts and material formulations for the present implementation are summarized in Table 1. The cervical ligaments were represented using discrete spring elements [34], and the occiput–C2 complex incorporated published rotational stiffness responses for flexion–extension and axial rotation [35].

**Table 1.** Model composition and material definitions for the APL Head-Neck model.

| Structure | Element type | Elements | Material model | LS-DYNA ID |
| --- | --- | --- | --- | --- |
| Anulus fibrosus | Hexahedral | 380 | Visco-soft-tissue | MAT_091 |
| Brain | Hexahedral | 246,300 | Bergström–Boyce rubber | MAT_269 |
| Bridging veins | Beam | 22 | Elastic | MAT_001 |
| Cerebrospinal fluid | Hexahedral | 90,900 | Viscoelastic | MAT_006 |
| Cortical bone | Quadrilateral | 45,300 | Elastic | MAT_001 |
| Dura mater | Quadrilateral | 18,400 | Elastic | MAT_001 |
| Falx–tentorium | Quadrilateral | 4,600 | Elastic–plastic–thermal | MAT_004 |
| Flesh | Hexahedral | 297,500 | Ogden rubber | MAT_077_O |
| Nucleus pulposus | Hexahedral | 510 | Mooney–Rivlin rubber | MAT_027 |
| Sinuses | Hexahedral | 22,600 | Elastic | MAT_001 |
| Skin | Quadrilateral | 28,100 | Elastic | MAT_001 |
| Trabecular bone | Hexahedral | 143,500 | Elastic | MAT_001 |
| Vertebral body | Hexahedral | 4,000 | Rigid | MAT_020 |

**Figure 1.**
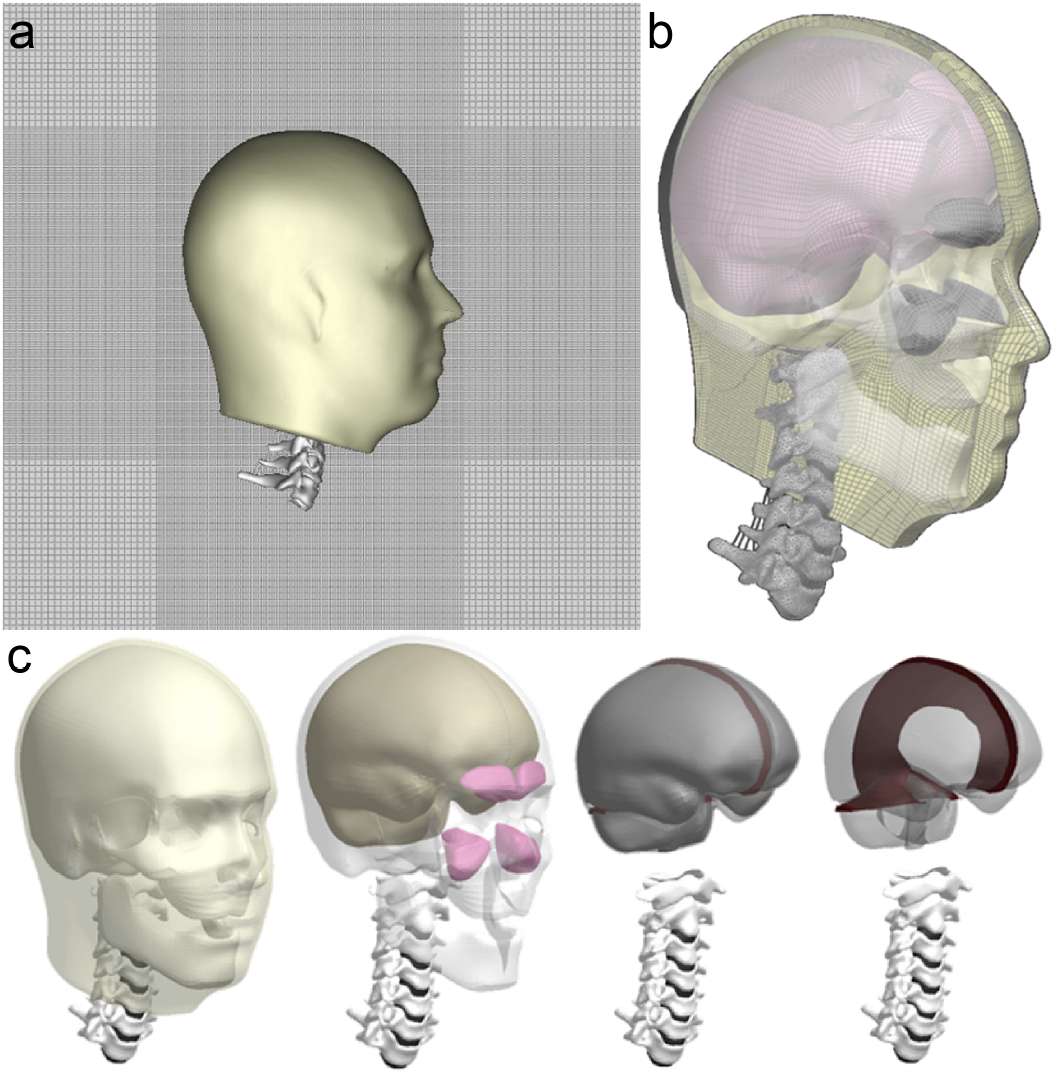
The ALE FE model used to simulate BOP propagation in the head. (a) The Lagrangian human head anatomy model positioned within a grid of Eulerian air domain elements shown in gray (500 × 500 × 500 mm). (b) The APL Head-Neck model includes cervical vertebrae, skull, and corresponding soft tissues. (c) Key anatomical structures include the flesh (tan), brain (pink), intervertebral disks (black), cervical vertebrae (white), cervical ligaments (gray), skull (white), sinuses (pink), and falx cerebri and tentorium cerebelli (red).

The selection and calibration of the brain constitutive model is essential to the modeling of BOP propagation within the brain. The brain was modeled using the Bergström–Boyce rubber model, implemented in LS-DYNA as MAT_269. The complete constitutive parameters and units are provided in Supplementary Table S1 [36, 37]. This material model has been previously applied to biomaterials and enables nonlinear generalization of viscoelastic materials that exhibit time-dependence and hysteresis under large deformations or in response to sudden loads [36]. Material characterization of brain tissue in tension, compression, and shear was previously conducted by Jin et al. [38]. Material parameters were previously determined through single element simulations of these experiments and are presented in Supplementary Table S1. This material model was previously validated under rotational loading using the response data published by Alshareef et al. [39]. Additionally, intracranial pressure (ICP) and shear strain responses were evaluated with respect to previous studies of the brain response to simulated brain blast exposures of Panzer et al. [29].

## Single blast exposure simulation

BOP exposures were selected based on the ranges measured during various military training exercises, such as explosive breaching and firing shoulder-mounted weapons, artillery, mortars, and .50-caliber guns. [8] During these training exercises, BOP exposures ranged from 4–90 kPa in peak pressure and from 1–90 kPa ms in peak impulse. To simulate the majority of these ranges, the exposures were selected to encompass the maximum possible impulse variation observed by Wiri (Figure 2) based on the recorded peak-pressure range (4–90 kPa) and positive-phase-duration range (2.2–10 ms). Intermediate pressures of 55.5 kPa, 21 kPa, and 4 kPa were selected at the same positive-phase duration as the maximum-impulse condition to evaluate the range of ICP and shear strains occurring under possible low-dose, short-duration conditions. Intermediate durations were selected at 2.2 ms, 6.1 ms, and 10 ms for the 21 kPa and 4 kPa conditions. Figure 2 shows a plot of the simulation conditions overlaid with previously published computational studies of blast traumatic brain injury (bTBI), as well as previously published blast injury thresholds for 50% lethality [40], pulmonary injury [41], and tympanic membrane rupture [42].

**Figure 2.**
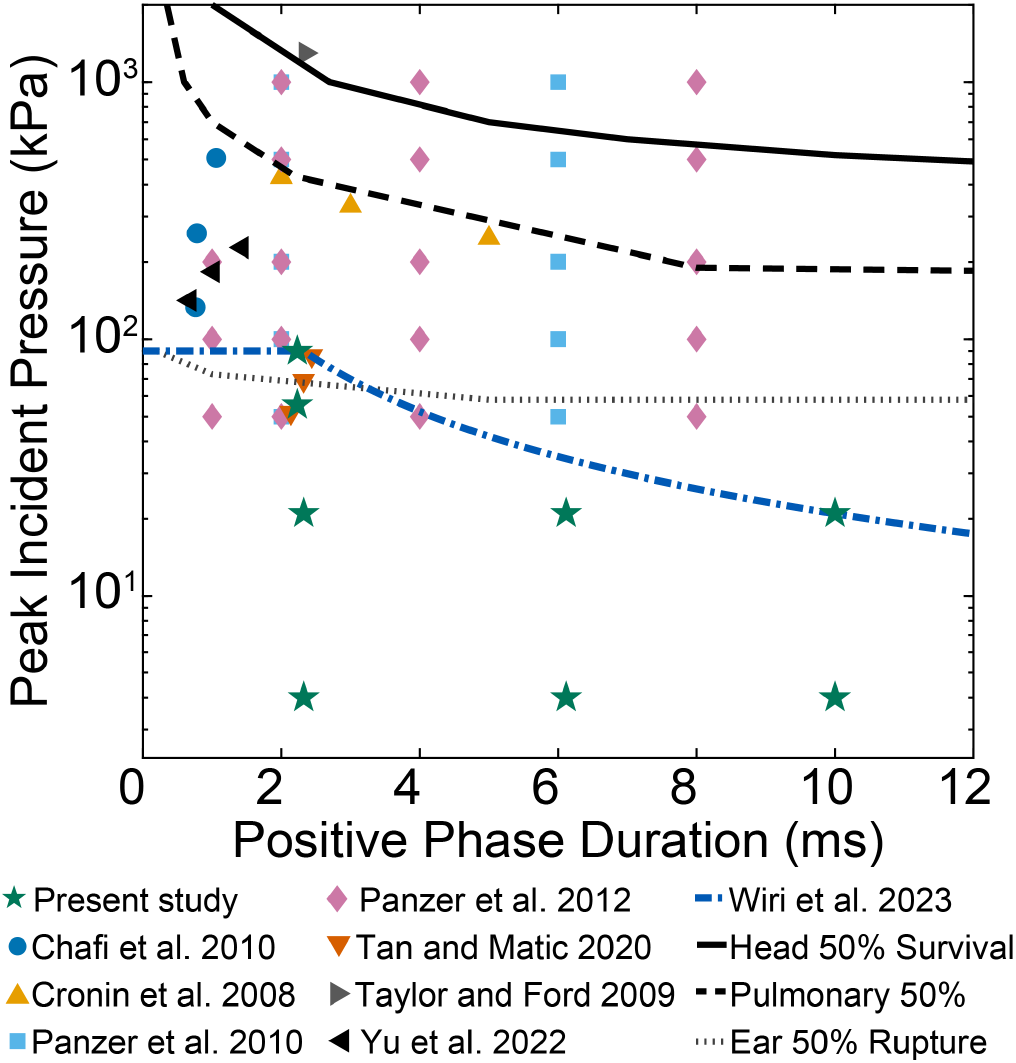
BOP simulation conditions used in this study, shown alongside conditions from previous computational bTBI studies. The envelope of peak pressures and impulses reported by Wiri et al. [8] was converted to equivalent peak-pressure and positive-phase-duration pairs (blue dash-dotted line), representing LIB exposures measured during military exercises. Previously published blast injury thresholds for 50% lethality [40], pulmonary injury [41], and tympanic membrane rupture [42] are also shown to provide context for higher-severity injury regimes.

The coupled ALE with load blast enhanced (LBE) method was selected to simulate the blast shock wave within LS-DYNA. This method models the explosive event based on the empirical model outlined in TM 5-855 US Army Technical Manual, which is implemented by CONWEP [43]. This includes modeling of the charge, explosive expansion after detonation, and the near-field physical effects. For air blast, it is efficient for single shock waves and significantly more efficient than modeling the charge in conditions where surface reflections can be assumed as minimally impacting on results [44]. The resulting pressures were then quantified at a distance where near-field effects are not substantial based on experimental data integrated into LS-DYNA. The nonlinearities of the shock wave propagation through the air domain were modeled using a polytropic equation of state to initialize at atmospheric pressure, 101.3 kPa. A 100 × 100 × 100 mesh was developed using the Structured-ALE (S-ALE) solver. The center of the blast was oriented to be in the same plane as the head center of gravity. After the impulse from the shock wave was deemed sufficiently transferred (at approximately 10 ms) the ALE domain was removed. Simulations were executed using LS-DYNA R14.0 double precision massively parallel processing (MPP) solver.

### Multiple blast exposure simulation

A second simulation experiment was conducted to characterize high-rate repetitive LIB exposure. As a representative worst-case scenario, the BOP from an M134 was simulated because its firing rate can reach 6,000 rounds per minute, corresponding to 10 ms between rounds [45]. Controlled simulation of multiple detonations was achieved by extracting the incident pressure developed in the previously described simulations for each of the 21 kPa peak BOP conditions with positive phase durations of 2.2 ms, 6.1 ms, and 10 ms. A peak BOP of 21 kPa was selected for its similarity to the average breacher exposure reported in Wiri et al. [46]. The extracted simulation pressures were applied as a boundary condition to the air domain within the S-ALE model using the load segment set approach within LS-DYNA. The incident pressure developed by the LBE definition in the previously described simulations was extracted for each of the 21 kPa conditions. The exposure interval was therefore set to 10 ms between BOP arrivals, corresponding to the maximum reported M134 firing rate. To evaluate accumulation of the ICP, three exposures were simulated in succession. The simulation was then run for 30 ms total using LS-DYNA.

### Simulation analysis

The ICP and shear strain were extracted both temporally and spatially from key elements within the model for the full duration of simulated BOP exposure using custom scripts in MATLAB 2023b (The MathWorks, Inc., Natick, MA, USA). A subset of elements (*n* = 1, 982) within the brain were randomly selected throughout the brain to measure blast pressure transmission to provide congruent analysis across simulations. These selected elements enabled the characterization of the entire brain tissue biomechanical response, rather than investigation of a localized region, while also preserving computational efficiency. To minimize the influence of a single-element response on the characterization of brain response, the presented corridors are provided as the 5th- and 95th-percentile peak values for each blast simulation condition. The ICP and the shear strain were analyzed for each simulated condition generated using CONWEP parameters in LS-DYNA. ICP results were similarly visualized in OASYS D3plot version 19.0 to qualitatively assess wave propagation through sections of interest within the brain [47].

Time-frequency analysis was conducted to characterize the frequency content of the ICP waveform with increasing exposure. A continuous wavelet transform (CWT) decomposition using a bump wavelet with 48 voices per octave was computed for all elements in MATLAB 2023b. The ICP signal was decomposed into a two-dimensional time-frequency representation, where the wavelet coefficients correspond to the similarity between the wavelet and the signal at different scales (inversely related to frequency) and temporal locations. The mean magnitude of the CWT across all elements was computed, quantifying the overall frequency content of the ICP signal. Results are reported using a scalogram generated in MATLAB.

## Results

### Simulated Single Blast Exposure Brain Response

Blast wave pressure transmission from air through flesh, skull, and into brain tissue is a complex and dynamic process. Prior to interaction with the human head surrogate, a planar wave was allowed to develop within the Eulerian air domain over approximately 0.3 ms. As shown in Figure 3, pressure begins to rise in the brain approximately 0.8 ms after the start of loading, which is defined here as time zero. The incident pressure wave then traveled anterior to posterior (coup to contrecoup) through the axial plane of the brain in less than 0.2 ms and started to develop a negative pressure reflection at the occiput. In parallel, the skull acted as a waveguide for the pressure wave, with a high positive pressure response moving along the skull on both sides anterior to posterior over 0.4 ms. Due to the higher acoustic impedance of the skull, as well as the falx cerebri and tentorium cerebelli, localized high-pressure regions began to form. These high-pressure regions then propagated along the skull-brain interface (1.0–1.3 ms after detonation). This positive pressure wave was followed by an area of negative pressure as the material responded to the initial loading. These positive and negative waves eventually reflected at the occiput around 0.8 ms after initial loading. Wave interference effects dominated the later response, resulting in the characteristic frequency of the ICP wave over time (1.4–1.7 ms after detonation and later).

**Figure 3.**
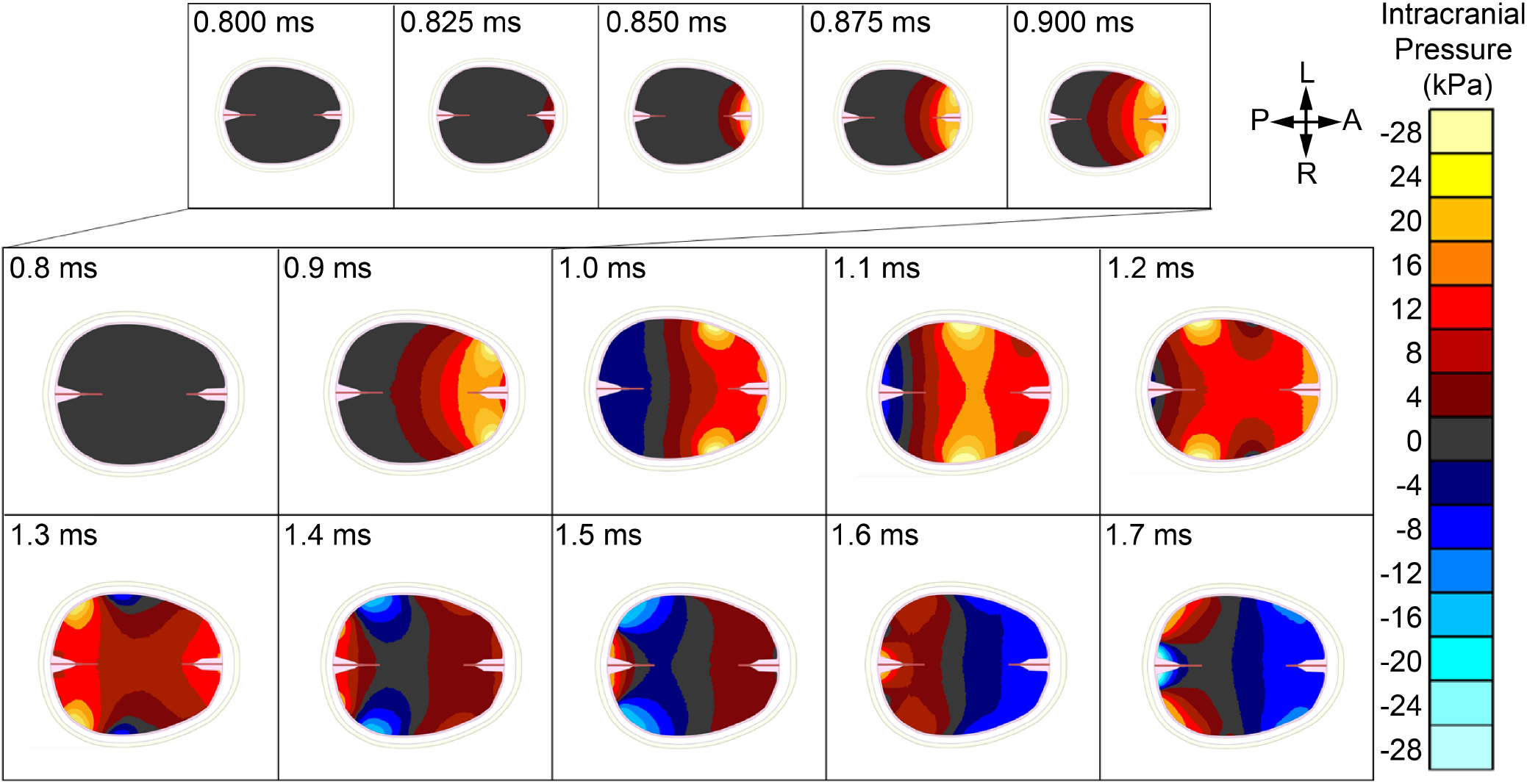
Example of the transient ICP observed during blast exposure, shown here for the 21 kPa incident blast overpressure with 6.1 ms duration. Anatomical orientation is indicated as posterior (P), anterior (A), left (L), and right (R).

To investigate this complex wave transmission through the brain in response to variation in the length of the incident wave’s positive phase, we compared the ICP of the 4 kPa simulations with positive-phase durations of 2.2 ms and 10 ms. Figure 4a presents the first 10 ms of transmission, where a slight phase shift can be observed between the two simulations, as well as differences in peak and minimum ICP responses. Analysis of brain response requires investigation of local responses in addition to the generalized response within the organ. By considering the local response at three distinct locations within the brain (Figure 4c), the coup and contrecoup responses can be studied, as well as comparisons between the medial and lateral responses. Positive pressure propagation initiated within the brain at 0.8 ms in the coup region (Figure 4c), and pressure peaked at the lateral element at 1.2 ms. As the wave propagated, negative pressure developed in the posterior region by 1.1 ms, creating tension in this focal area.

**Figure 4.**
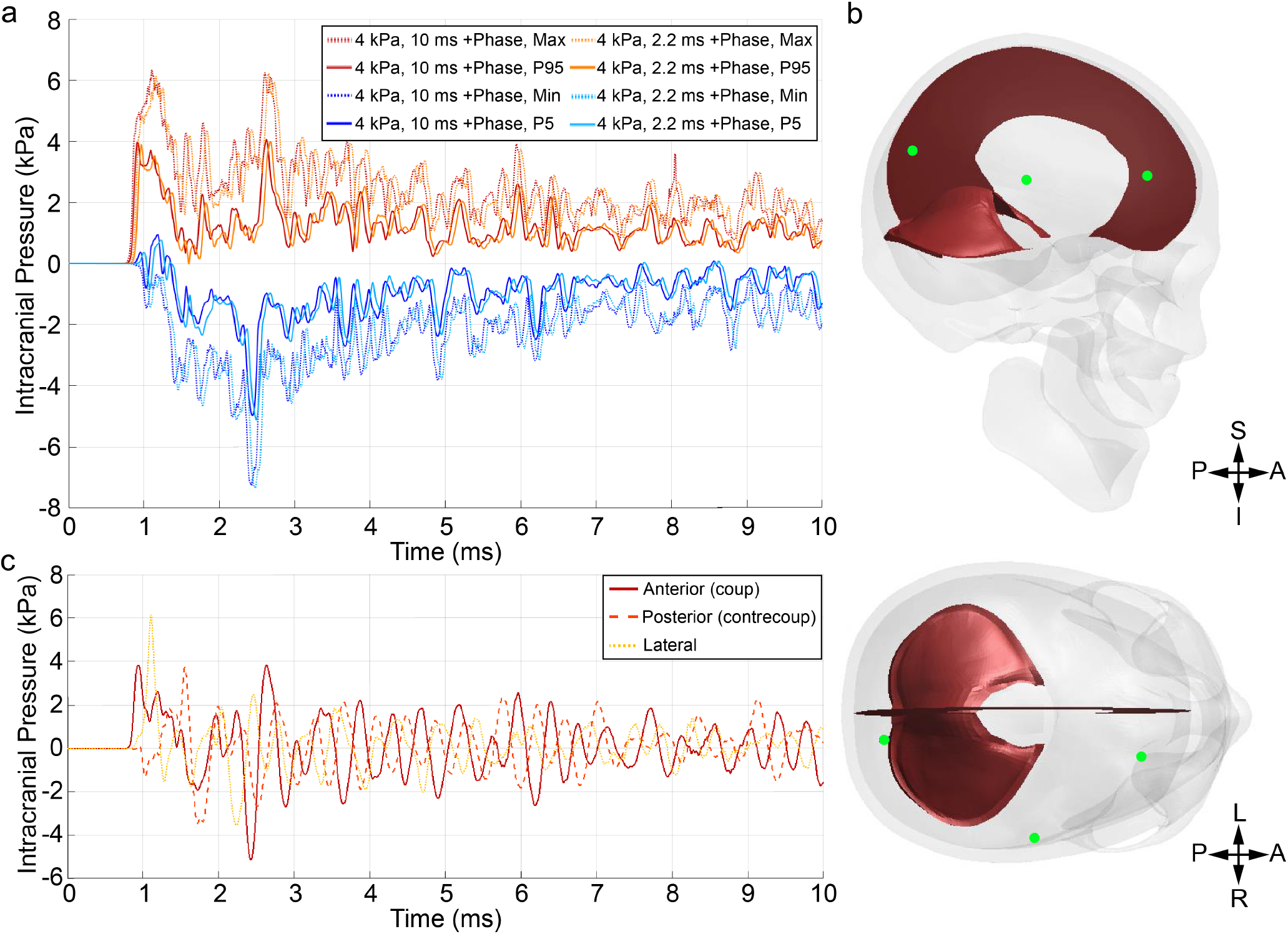
Transient ICP responses to Friedlander waves with a peak incident pressure of 4 kPa. (a) Minimum, 5th percentile (P5), 95th percentile (P95), and maximum ICP histories for positive-phase durations of 2.2 and 10 ms. (b) Sagittal view of the three sampled brain elements (green). (c) ICP histories at the anterior (coup), posterior (contrecoup), and lateral elements, accompanied by an axial view of their locations. Anatomical orientation is indicated as posterior (P), anterior (A), superior (S), inferior (I), left (L), and right (R).

The dynamic brain response after blast exposure was quantified using ICP and the shear strain induced by the tension and compression of biological tissue. These metrics provide insight into the injurious potential of blast overpressure (BOP) loading on brain tissue structure and function. Peak values for both metrics are presented in Table 2, along with times to peak for each of the simulated conditions. Comparing the visualized response for the 21 kPa incident blast overpressure with 6.1 ms duration condition shown in Figure 3 to the results in Table 2, the peak ICP occurred during this initial loading phase and not during the reflections. Note that the *Time of Peak* indicated in Table 2 is similar to that in Figure 3 in that the timing starts at the start of simulation, not at the start of the initial loading of the head. As such, for the 21 kPa incident blast overpressure, the peak ICP occurred around 0.9 ms or in the second image of the top row in Figure 3. The timing of the peak ICP trended linearly with peak incident pressure for the first three conditions, becoming dramatically larger at the lowest peak incident pressure. The recorded peak ICP, however, did scale linearly with peak incident pressure. The maximum ICP exceeded the peak incident BOP due to observed interactions between compressive pressure waves entering from the anterior skull and from the lateral skull surfaces, resulting in constructive interference and higher ICP than the anticipated attenuated incident BOP [48]. Peak ICP varied little with positive-phase duration at a fixed incident pressure. At 21 kPa, peak shear strain was greatest for the 10.0-ms positive phase, and time to peak was longer for the shorter-duration conditions. At 4 kPa, differences in peak shear strain were below the reported precision, and the peak was not reached within 15 ms.

**Table 2.**
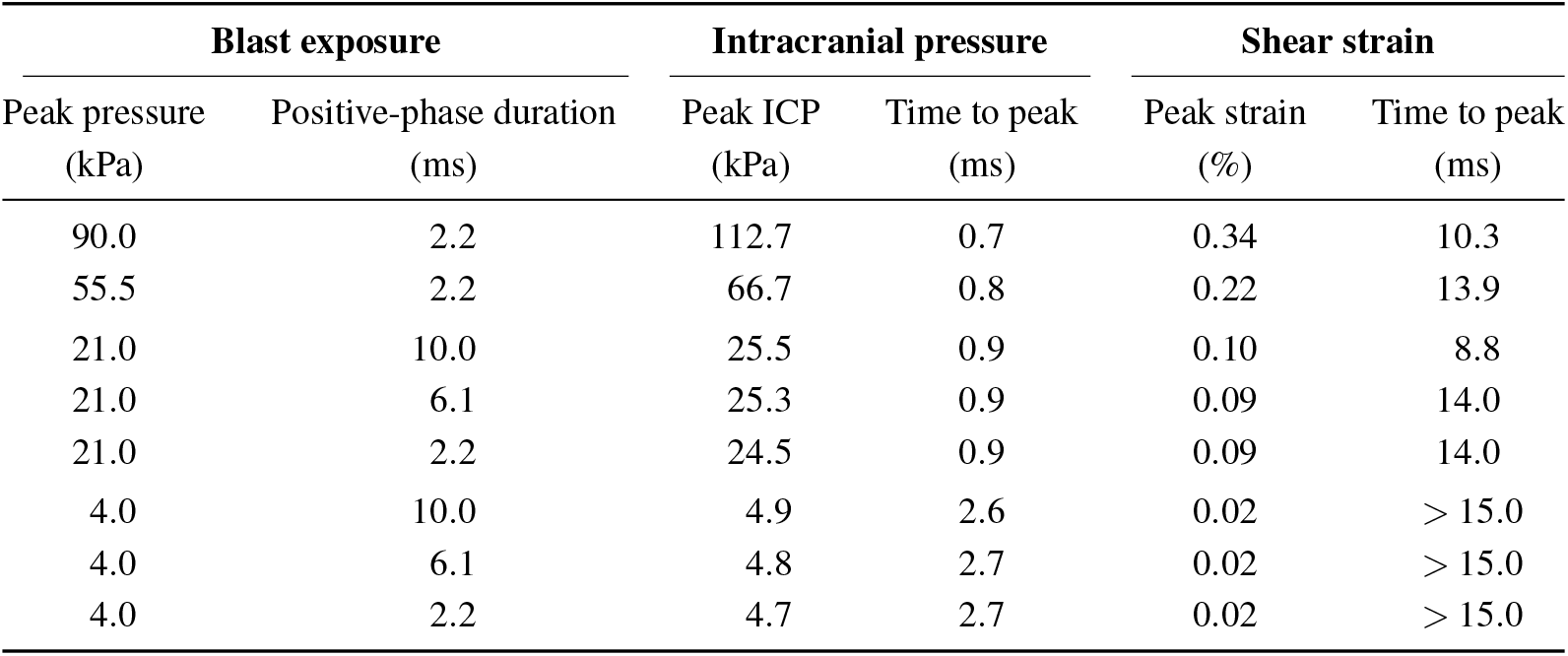
Peak ICP and shear-strain responses at the 95th percentile across sampled brain elements. Blast exposure conditions are nominal simulation inputs. Values greater than 15.0 ms indicate that the peak shear strain was not reached within the analysis window.

| Blast exposure |  | Intracranial pressure |  | Shear strain |  |
| --- | --- | --- | --- | --- | --- |
| Peak pressure<br>(kPa) | Positive-phase duration<br>(ms) | Peak ICP<br>(kPa) | Time to peak<br>(ms) | Peak strain<br>(%) | Time to peak<br>(ms) |
| 90.0 | 2.2 | 112.7 | 0.7 | 0.34 | 10.3 |
| 55.5 | 2.2 | 66.7 | 0.8 | 0.22 | 13.9 |
| 21.0 | 10.0 | 25.5 | 0.9 | 0.10 | 8.8 |
| 21.0 | 6.1 | 25.3 | 0.9 | 0.09 | 14.0 |
| 21.0 | 2.2 | 24.5 | 0.9 | 0.09 | 14.0 |
| 4.0 | 10.0 | 4.9 | 2.6 | 0.02 | > 15.0 |
| 4.0 | 6.1 | 4.8 | 2.7 | 0.02 | > 15.0 |
| 4.0 | 2.2 | 4.7 | 2.7 | 0.02 | > 15.0 |

### Simulated Multiple Blast Exposure Brain Response

Figure 5 illustrates the 5th and 95th percentile pressures predicted within the brain during a single simulation involving three consecutive 21 kPa incident BOP exposures. Panels 5a, 5c, and 5e present ICP results for simulations with BOP wave durations of 2.2, 6.1, and 10 ms, respectively. The simulated peak ICP increased with successive BOP exposures regardless of duration, with the magnitude of ICP at the third BOP exposure showing the greatest increase when exposed to the longest duration BOP. This study introduces a methodology for evaluating the accumulation of mechanical stresses within the brain during repeated pressure-wave exposures. The ICP responses show increasing spectral amplitudes in both the 5th- and 95th-percentile signals with each successive blast exposure. Panels 5b, 5d, and 5f illustrate changes in spectral content across successive exposures with identical BOP magnitude and duration. For reference, the tail of each white arrow indicates the onset of the BOP, while the arrow length indicates its duration. The arrival of each pressure wave is characterized by high-frequency content (*>*2 kHz), whose amplitude increases with each successive exposure within a given simulation. During the initial exposure, the amplitude of this high-frequency content also increases with BOP duration. After approximately the first 2 ms, pressure-wave reflections within the cranial cavity and brain tissue produce lower-frequency content (*<*2 kHz). This content is concentrated near the characteristic resonant frequencies of the coupled brain–cranium system and increases in amplitude with both incident BOP duration and the number of successive exposures.

**Figure 5.**
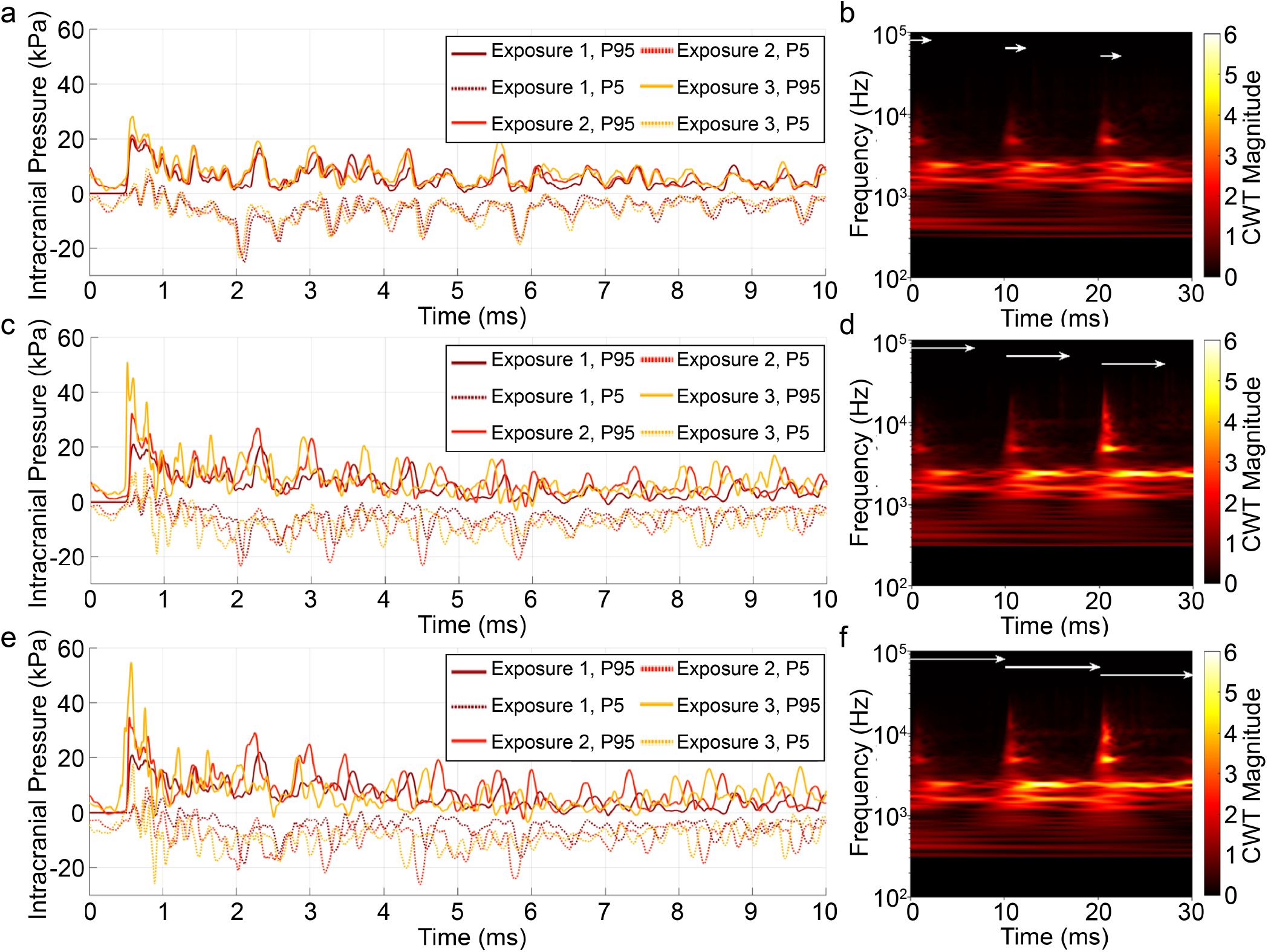
5th percentile (P5) and 95th percentile (P95) ICP responses and spectral content for three detonations at 10-ms intervals. Three Friedlander waves were simulated with a peak incident pressure of 21 kPa. Panels (a), (c), and (e) show ICP responses for positive-phase durations of 2.2, 6.1, and 10 ms, respectively; panels (b), (d), and (f) show the corresponding continuous wavelet transform (CWT) coefficient magnitudes.

## Discussion

Due to the complexities of *in vivo* brain responses to traumatic injury, computational approaches have been applied to study both direct and indirect injury mechanisms of TBI. However, most studies focus on single, higher-dose events, leaving a gap in understanding LIB conditions commonly observed in military training environments. Through refinement and application of a validated human head-brain FE model, we characterized the LIB regime in the anterior-to-posterior loading direction, with peak pressures ranging from 4–90 kPa as observed in military training environments [8] and associated with adverse neurological outcomes including mbTBI [4, 11–13]. In this study, we quantified maximum ICP (Fig. 3), spectral magnitude (Fig. 5), and shear strains (Table 2) under single and repetitive LIB exposures. These metrics provide a consistent basis to compare single and repeated LIB exposures and to assess the relative importance of pressure, spectral content, and strain in potential injury mechanisms.

The simulations demonstrated that LIB produced localized regions of high positive pressure (*>*30 kPa) and high-magnitude negative pressure (*<*-30 kPa). ICP at the anterior skull surface was greater than that observed at the posterior skull surface, with the most lateral surface presenting the largest ICP at the skull curvature. For all incident pressures and positive phase durations reported in Table 2, the simulated peak ICP exceeded the incident pressure, producing localized pressure amplifications that were consistent with prior simulation studies. Within this study, we predicted brain ICP values of 4.7–112.7 kPa. For example, Panzer et al. reported 65–72 kPa 95th percentile ICP for a 50 kPa incident overpressure with 1–8 ms duration [29]. Taylor et al. also observed elevated pressures and localized deviatoric stresses under blast loading [49]. The localized pressure amplifications observed here occurred during both initial propagation and intracranial reflection (Fig. 3), consistent with prior studies indicating that blast loading produces localized injury regions rather than uniform loading of the brain [50, 51]. For anterior blast exposure, the highest pressures occurred near the frontal cortex, while negative pressures developed near posterior surfaces [50]. This anterior-to-posterior pressure transmission agrees with experimental observations of blast wave propagation through the brain [25, 52]. These responses are typical of planar blast exposure, though occupational loading environments may also include reflections and confined geometries that amplify or reshape intracranial responses and should be considered when contextualizing field measurements [30].

As shown previously, skull curvature engaged during blast exposure strongly influences the magnitude and spatial extent of these pressure amplifications, with lateral curvature producing larger affected regions and more prominent intracranial pressure amplifications than frontal exposure alone [30]. Prior studies have shown that acoustic-impedance mismatches among the skull, cerebrospinal fluid, and brain can produce complex transmission and reflection of blast-induced pressure waves, resulting in spatially nonuniform intracranial loading [53, 54]. The falx and tentorium may further influence wave propagation by reflecting and focusing mechanical waves within the brain [55]. These responses are further influenced by the rate and duration of applied load, consistent with previous studies of brain tissue response under dynamic loading [56]. Some limitations should be considered when interpreting these results. The magnitude and spatial distribution of the ICP depend on anatomical variation, including differences in skull geometry (e.g., curvature, cortical and diploë thickness, and sinus morphology) [30, 57]. As a single model cannot capture the full range of anatomical variability, future work could explore sensitivity to both anatomical detail and material properties. Additionally, the relative reduction in peak ICP compared with simplified plane-strain models likely reflects the inclusion of the full 3D head geometry and blast aerodynamics, which introduce additional compliance and attenuation compared with constrained models [29]. These findings nevertheless suggest that specific brain regions may be disproportionately influenced by LIB exposure.

Although the 21 kPa incident BOP used in the repetitive-exposure simulations was below the DoD’s interim 28 kPa (4 psi) blast-overpressure safety threshold, the simulations showed that rapid repeated exposures resulted in accumulation of ICP and altered spectral content. [7] In the representative repetitive-loading case of three consecutive 21 kPa exposures separated by 10 ms, peak ICP increased by 8–26% relative to single exposures, with the greatest amplification by the third exposure and under the longest positive phase duration. The arrival of each pressure wave was marked by high-frequency content (*>*2 kHz) that intensified with successive exposures, whereas after approximately 2 ms intracranial reflections produced lower-frequency reverberations (*<*2 kHz) of increasing magnitude with both longer positive phase duration and repeated exposure. This response is likely driven by incomplete mechanical relaxation of brain tissue between exposures, resulting in re-exposure before full dissipation of energy occurs. These findings suggest a potential injury mechanism associated with the accumulation of biomechanical loading during repetitive, low-amplitude blast exposures, extending prior computational and experimental studies that focused on single, higher-pressure regimes. High-frequency components of blast pressure waves have been hypothesized to influence tissue injury; however, their biological relevance to brain tissue and any associated injury thresholds remain uncertain [58]. Accordingly, the increasing high-frequency content observed with repeated exposure warrants further investigation, even when peak pressures remain relatively small. Further computational and experimental studies of repeated exposure are necessary to better understand these phenomena and their relationship to injury. Such studies may define the dissipation timeline of intracranial stresses; however, in the present context shot cadence should be carefully considered to minimize accumulation of pressure and high-frequency pressure content within the brain.

Despite the complex pressure dynamics observed, predicted shear strains remained relatively small. In this study we predicted 95th percentile shear strains of approximately 0.02–0.34%. For the comparable condition of 55.5 kPa and 2.2 ms, the model predicted 66.7 kPa 95th percentile ICP and 0.22% 95th percentile shear strain, aligning well with the range reported by Panzer et al. (65–72 kPa ICP and 0.07–0.3% strain) under similar blast loading [29]. These values are several orders of magnitude lower than strain levels typically associated with inertial or blunt TBI, where rapid head accelerations cause large axonal stretching leading to diffuse axonal injury. In contrast, LIB is dominated by rapid pressure wave transmission through the skull and brain with comparatively small strains. This behavior is consistent with the high bulk modulus of brain tissue and the largely incompressible boundary imposed by the skull, which favor rapid pressure transmission over substantial shear deformation under LIB conditions, where minimal head acceleration is expected. As a result, the intracranial response is dominated by pressure wave dynamics, where the transmitted pressure wave undergoes multiple reflections within the intracranial cavity due to acoustic impedance mismatches between the cranium and brain [49]. These reflections produce oscillatory ICP responses with frequency content spanning approximately 0.5–10 kHz and persisting for roughly 1–10 ms [59]. While similar pressure transmission occurs in higher-severity blast exposures, the associated blast wind can induce substantial head acceleration, introducing additional strain-based injury mechanisms.

Even when global strain magnitudes remain small, localized pressure-time gradients can remain large, particularly during repeated exposures. Nonlinear viscoelastic propagation within brain tissue may further steepen these wave fronts and sustain high spatial and temporal gradients [53]. For example, predicted strains for a representative 21 kPa exposure, similar to average breacher exposures, were approximately 0.08–0.10%, while exposures near 4 kPa produced strains near 0.02% [8]. These values remain far below commonly cited diffuse axonal injury thresholds of approximately 15% strain [60]. Nevertheless, nanoscale abnormalities have been observed in animal studies following low-intensity blast exposure, [61] suggesting that alternative injury pathways may be involved. Additionally, in the case of repeated exposures, there may be accumulation of axonal damage, similar to those observed in vitro after repeated stretching. Future analyses may therefore benefit from examining strain-rate distributions and pressure gradients in addition to strain magnitude alone.

Several injury mechanisms have been proposed to contribute to mbTBI, including both direct mechanisms (e.g., acoustic impedance effects during wave propagation, cavitation phenomena, skull flexure–driven strain) and indirect mechanisms such as relative brain motion or thoracic coupling [54, 57, 62]. Characterizing which mechanisms dominate under LIB conditions remains an open challenge. In this study we focused on intracranial pressure propagation and shear strain as mechanistically relevant responses. The simulations demonstrated ICP oscillations and localized regions of positive and negative pressure resulting from wave reflections within the cranial cavity [49, 50, 63]. These oscillations produce transient tension phases within the brain that could hypothetically induce CSF cavitation. However, the relatively low pressures observed here suggest that cavitation is unlikely to dominate under the LIB conditions considered [29]. Instead, the results support the hypothesis that repeated pressure loading and high-frequency stress gradients may contribute to injury through cumulative biomechanical effects.

Within this study, a simplified head-brain geometry was deployed, consisting of the brain, falx and tentorium, skull cortex and diploë, CSF, dura, sinuses, and surrounding soft tissues. Previous studies have incorporated additional anatomical detail, including separation of gray and white matter or multiscale modeling approaches capturing axonal deformation [29, 64]. Such approaches may provide additional insight into localized injury mechanisms where high-frequency pressures exist. The predicted biomechanical response is also sensitive to the constitutive model used to represent brain tissue. Previous blast simulations have employed hyperelastic, viscoelastic, and hyperviscoelastic formulations to capture the rate- and frequency-dependent behavior of neural tissue [59, 65, 66]. In this study, the brain was modeled using a Bergström–Boyce constitutive formulation calibrated to experimental data [38]. While this model captures key nonlinear material behavior, unresolved microscale failure mechanisms remain beyond the resolution of continuum models. Additionally, because this study characterized only anterior-to-posterior loading, skull curvature may alter the response under lateral or posterior loading; additional studies are needed to characterize these directional effects. Finally, although the model incorporates the cervical spine, whole-body coupling and active musculature may influence intracranial strain responses. More detailed musculoskeletal coupling may therefore be required to fully characterize head–body interactions during blast exposure. Despite these limitations, the results provide a foundation for future multiscale computational and experimental studies aimed at clarifying the mechanisms of mbTBI under occupational blast exposures.

## Conclusion

An increasing body of evidence implicates low-magnitude repetitive blast overpressure exposures in long-term neurological complications among military personnel. Because neurological effects may develop gradually or remain subclinical after exposure, the mechanisms underlying brain injury as a result of LIB remain poorly understood. This study was among the first to computationally characterize the pressure, strain, and spectral content of the brain in response to low-magnitude blast overpressure exposures associated with Warfighter weaponry. These results provide insight into the need for low-dose blast research, demonstrate consistency with prior computational studies, and emphasize the importance of investigating repeated exposures to clarify injury mechanisms and cumulative dose effects.

As the study of human injury mitigation continues, animal, cadaveric, surrogate, and computational approaches remain key to developing our understanding of the intracranial brain biomechanical response. Of these, computational approaches offer the opportunity to ground proposed injury mechanism hypotheses in our understanding of biomechanical theory within a near-perfectly controlled environment, as well as evaluate new conceptual ideas efficiently. This and related efforts may also guide the design of future experimental studies of mbTBI by helping identify optimal repetitive LIB test conditions and providing a theoretical framework for test-procedure development and results analysis.

As future studies continue to investigate the neurological effects of mbTBI, in order to further our understanding of adverse neurological outcomes and identify intervention strategies, innovative strategies to study these complex phenomena will be paramount to success. Given the critical importance of establishing occupational blast exposure limits during warfighting activities, continued study of inter-shot intervals is needed to characterize intracranial energy dissipation. These responses are likely to be sensitive to headborne equipment, posture during exposure, individual anthropometry, and loading direction; these factors should be carefully considered during experimental study design. Validation of these computational studies requires the development of high-quality, representative experimental datasets, such as those presented by Wiri et al., with instrumentation of training participants to capture repetitive blast exposure dose across multiple exposures. Computationally, incorporation of multiscale modeling could enable study of microscale failure models and greater granularity of tissue response. In turn, this would enable more robust consideration of pressure gradients and strain rates relevant to biomechanical injury hypotheses. Until effective mitigation strategies become available for military personnel who face repetitive blast exposure, the approaches outlined here can continue to inform efforts to reduce the risk of mbTBI and its symptoms.

## Data availability

The data supporting the findings of this study are available from the corresponding author upon reasonable request.

## Acknowledgements

This work was supported by the Johns Hopkins University Applied Physics Laboratory Research and Exploratory Development Department through the Independent Research and Development Program.

## Author contributions statement

Conceptualization: V.D.A., E.B.-K., M.D.Y., and T.A.M.; Investigation: M.D.Y.; Formal analysis: M.D.Y., T.A.M., and E.B.-K.; Writing–review & editing: M.D.Y., T.A.M., V.D.A., K.A.O., and E.B.-K.

## References

1. Kamimori, G. H., Reilly, L. A., LaValle, C. R. & Olaghere Da Silva, U. B. Occupational overpressure exposure of breachers and military personnel. Shock. Waves 27, 837–847, DOI: 10.1007/s00193-017-0738-4 (2017).

2. Stocker, R. P. et al. Combat-related blast exposure and traumatic brain injury influence brain glucose metabolism during REM sleep in military veterans. NeuroImage 99, 207–214, DOI: 10.1016/j.neuroimage.2014.05.067 (2014).

3. Thangavelu, B. et al. Overpressure Exposure From .50-Caliber Rifle Training Is Associated With Increased Amyloid Beta Peptides in Serum. Front. Neurol. 11, 620, DOI: 10.3389/fneur.2020.00620 (2020).

4. Belding, J. N. et al. Potential Health and Performance Effects of High-Level and Low-Level Blast: A Scoping Review of Two Decades of Research. Front. Neurol. 12, 628782, DOI: 10.3389/fneur.2021.628782 (2021).

5. Boutté, A. M. et al. Neurotrauma Biomarker Levels and Adverse Symptoms Among Military and Law Enforcement Personnel Exposed to Occupational Overpressure Without Diagnosed Traumatic Brain Injury. JAMA Netw. Open 4, e216445, DOI: 10.1001/jamanetworkopen.2021.6445 (2021).

6. Gilmore, N. et al. Impact of repeated blast exposure on active-duty United States Special Operations Forces. Proc. Natl. Acad. Sci. 121, e2313568121, DOI: 10.1073/pnas.2313568121 (2024).

7. Hicks, K. H. Department of Defense Requirements for Managing Brain Health Risks from Blast Overpressure. Tech. Rep., U.S. Department of Defense (2024).

8. Wiri, S. et al. Dynamic monitoring of service members to quantify blast exposure levels during combat training using BlackBox Biometrics Blast Gauges: explosive breaching, shoulder-fired weapons, artillery, mortars, and 0.50 caliber guns. Front. Neurol. 14, 1175671, DOI: 10.3389/fneur.2023.1175671 (2023).

9. Rosenfeld, J. V. et al. Blast-related traumatic brain injury. The Lancet Neurol. 12, 882–893, DOI: 10.1016/S1474-4422(13)70161-3 (2013).

10. Siedhoff, H. R. et al. Perspectives on Primary Blast Injury of the Brain: Translational Insights Into Non-inertial Low-Intensity Blast Injury. Front. Neurol. 12, 818169, DOI: 10.3389/fneur.2021.818169 (2022).

11. Carr, W. et al. Relation of Repeated Low-Level Blast Exposure With Symptomology Similar to Concussion. J. Head Trauma Rehabil. 30, 47–55, DOI: 10.1097/HTR.0000000000000064 (2015).

12. Sajja, V. S. S. S. et al. The Role of Very Low Level Blast Overpressure in Symptomatology. Front. Neurol. 10, 891, DOI: 10.3389/fneur.2019.00891 (2019).

13. Tate, C. M. et al. Serum Brain Biomarker Level, Neurocognitive Performance, and Self-Reported Symptom Changes in Soldiers Repeatedly Exposed to Low-Level Blast: A Breacher Pilot Study. J. Neurotrauma 30, 1620–1630, DOI: 10.1089/neu.2012.2683 (2013).

14. Trotter, B. B., Robinson, M. E., Milberg, W. P., McGlinchey, R. E. & Salat, D. H. Military blast exposure, ageing and white matter integrity. Brain 138, 2278–2292, DOI: 10.1093/brain/awv139 (2015).

15. McKee, A. C. & Robinson, M. E. Military-related traumatic brain injury and neurodegeneration. Alzheimer’s & Dementia 10, DOI: 10.1016/j.jalz.2014.04.003 (2014).

16. Barnes, D. E. et al. Association of Mild Traumatic Brain Injury With and Without Loss of Consciousness With Dementia in US Military Veterans. JAMA Neurol. 75, 1055, DOI: 10.1001/jamaneurol.2018.0815 (2018).

17. Taber, K. H. et al. White Matter Compromise in Veterans Exposed to Primary Blast Forces. J. Head Trauma Rehabil. 30, E15–E25, DOI: 10.1097/HTR.0000000000000030 (2015).

18. Mac Donald, C. L. et al. Detection of Blast-Related Traumatic Brain Injury in U.S. Military Personnel. New Engl. J. Medicine 364, 2091–2100, DOI: 10.1056/NEJMoa1008069 (2011).

19. Jorge, R. E. et al. White Matter Abnormalities in Veterans With Mild Traumatic Brain Injury. Am. J. Psychiatry 169, 1284–1291, DOI: 10.1176/appi.ajp.2012.12050600 (2012).

20. Olivera, A. et al. Peripheral Total Tau in Military Personnel Who Sustain Traumatic Brain Injuries During Deployment. JAMA Neurol. 72, 1109, DOI: 10.1001/jamaneurol.2015.1383 (2015).

21. Pattinson, C. L. et al. Elevated Tau in Military Personnel Relates to Chronic Symptoms Following Traumatic Brain Injury. J. Head Trauma Rehabil. 35, 66–73, DOI: 10.1097/HTR.0000000000000485 (2020).

22. Leonardi, A. D. C. et al. Methodology and Evaluation of Intracranial Pressure Response in Rats Exposed to Complex Shock Waves. Annals Biomed. Eng. 41, 2488–2500, DOI: 10.1007/s10439-013-0850-2 (2013).

23. Chavko, M., Koller, W. A., Prusaczyk, W. K. & McCarron, R. M. Measurement of blast wave by a miniature fiber optic pressure transducer in the rat brain. J. Neurosci. Methods 159, 277–281, DOI: 10.1016/j.jneumeth.2006.07.018 (2007).

24. Feng, K. et al. Biomechanical Responses of the Brain in Swine Subject to Free-Field Blasts. Front. Neurol. 7, DOI: 10.3389/fneur.2016.00179 (2016).

25. Iwaskiw, A. S. et al. The measurement of intracranial pressure and brain displacement due to short-duration dynamic overpressure loading. Shock. Waves 28, 63–83, DOI: 10.1007/s00193-017-0759-z (2018).

26. Norris, C. et al. Bilayer surrogate brain response under various blast loading conditions. Shock. Waves 34, 357–367, DOI: 10.1007/s00193-024-01158-5 (2024).

27. Wermer, A. et al. Materials Characterization of Cranial Simulants for Blast-Induced Traumatic Brain Injury. Mil. Medicine 185, 205–213, DOI: 10.1093/milmed/usz228 (2020).

28. Knutsen, A. K. et al. Characterization of material properties and deformation in the ANGUS phantom during mild head impacts using MRI. J. Mech. Behav. Biomed. Mater. 138, 105586, DOI: 10.1016/j.jmbbm.2022.105586 (2023).

29. Panzer, M. B., Myers, B. S., Capehart, B. P. & Bass, C. R. Development of a Finite Element Model for Blast Brain Injury and the Effects of CSF Cavitation. Annals Biomed. Eng. 40, 1530–1544, DOI: 10.1007/s10439-012-0519-2 (2012).

30. Tan, X. G. & Matic, P. Simulation of Cumulative Exposure Statistics for Blast Pressure Transmission Into the Brain. Mil. Medicine 185, 214–226, DOI: 10.1093/milmed/usz308 (2020).

31. Yates, M. et al. Computational Assessment of Headborne Equipment: Alteration of Head and Neck Biomechanics During Blast-Induced Accelerative Loading. In 16th Personal Armour Systems Symposium (PASS. 2023), 174–184, DOI: 10.52202/080042-0019 (Royal Military Academy (Belgium), Dresden, Germany, 2023).

32. Gordon, C. C. et al. 2012 Anthropometric Survey of U.S. Army Personnel: Methods and Summary Statistics. Tech. Rep. NATICK/TR-15/007, U.S. Army Natick Soldier Research, Development and Engineering Center, Natick, MA (2014).

33. Ackerman, M. J. The Visible Human Project: a resource for education. Acad. Medicine 74, 667–70, DOI: 10.1097/00001888-199906000-00012 (1999).

34. Brolin, K. & Halldin, P. Development of a Finite Element Model of the Upper Cervical Spine and a Parameter Study of Ligament Characteristics:. Spine 29, 376–385, DOI: 10.1097/01.BRS.0000090820.99182.2D (2004).

35. Panjabi, M. M. et al. Development of a System for In Vitro Neck Muscle Force Replication in Whole Cervical Spine Experiments:. Spine 26, 2214–2219, DOI: 10.1097/00007632-200110150-00012 (2001).

36. Bergström, J. & Boyce, M. Constitutive modeling of the time-dependent and cyclic loading of elastomers and application to soft biological tissues. Mech. Mater. 33, 523–530, DOI: 10.1016/S0167-6636(01)00070-9 (2001).

37. Dal, H. & Kaliske, M. Bergström–Boyce model for nonlinear finite rubber viscoelasticity: theoretical aspects and algorithmic treatment for the FE method. Comput. Mech. 44, 809–823, DOI: 10.1007/s00466-009-0407-2 (2009).

38. Jin, X., Zhu, F., Mao, H., Shen, M. & Yang, K. H. A comprehensive experimental study on material properties of human brain tissue. J. Biomech. 46, 2795–2801, DOI: 10.1016/j.jbiomech.2013.09.001 (2013).

39. Alshareef, A., Wu, T., Giudice, J. S. & Panzer, M. B. Toward subject-specific evaluation: methods of evaluating finite element brain models using experimental high-rate rotational brain motion. Biomech. Model. Mechanobiol. 20, 2301–2317, DOI: 10.1007/s10237-021-01508-7 (2021).

40. Rafaels, K. et al. Survival Risk Assessment for Primary Blast Exposures to the Head. J. Neurotrauma 28, 2319–2328, DOI: 10.1089/neu.2009.1207 (2011).

41. Bass, C. R., Rafaels, K. A. & Salzar, R. S. Pulmonary Injury Risk Assessment for Short-Duration Blasts. J. Trauma: Inj. Infect. & Critical Care 65, 604–615, DOI: 10.1097/TA.0b013e3181454ab4 (2008).

42. Richmond, D. R., Fletcher, E. R., Yelverton, J. T. & Phillips, Y. Y. Physical Correlates of Eardrum Rupture. Annals Otol. Rhinol. & Laryngol. 98, 35–41, DOI: 10.1177/00034894890980S507 (1989).

43. Hyde, D. W. Microcomputer Programs CONWEP and FUNPRO, Applications of TM 5-855-1, “Fundamentals of Protective Design for Conventional Weapons” (User’s Guide). Tech. Rep. SL-88-1, U.S. Army Engineer Waterways Experiment Station, Structures Laboratory, Vicksburg, MS (1988).

44. Abedini, M., Zhang, C., Mehrmashhadi, J. & Akhlaghi, E. Comparison of ALE, LBE and pressure time history methods to evaluate extreme loading effects in RC column. Structures 28, 456–466, DOI: 10.1016/j.istruc.2020.08.084 (2020).

45. Bureau of Alcohol, Tobacco, Firearms and Explosives. ATF ruling 2004-5: Mini-gun ruling. ATF Ruling 2004-5, U.S. Department of Justice (2004).

46. Wiri, S., Ritter, A. C., Bailie, J. M., Needham, C. & Duckworth, J. L. Computational modeling of blast exposure associated with recoilless weapons combat training. Shock. Waves 27, 849–862, DOI: 10.1007/s00193-017-0755-3 (2017).

47. Oasys Limited. Oasys Suite 19.0 (2022).

48. Sundaramurthy, A. Understanding the Effects of Blast Wave on the Intracranial Pressure and Traumatic Brain Injury in Rodents and Humans Using Experimental Shock Tube and Numerical Simulations. Ph.D. thesis, University of Nebraska– Lincoln Lincoln, NE (2014).

49. Taylor, P. A. & Ford, C. C. Simulation of Blast-Induced Early-Time Intracranial Wave Physics leading to Traumatic Brain Injury. J. Biomech. Eng. 131, 061007, DOI: 10.1115/1.3118765 (2009).

50. Wang, C. et al. Computational Study of Human Head Response to Primary Blast Waves of Five Levels from Three Directions. PLoS ONE 9, e113264, DOI: 10.1371/journal.pone.0113264 (2014).

51. Rezaei, A., Salimi Jazi, M. & Karami, G. Computational modeling of human head under blast in confined and open spaces: Primary blast injury. Int. J. for Numer. Methods Biomed. Eng. 30, 69–82, DOI: 10.1002/cnm.2590 (2014).

52. Salzar, R. S., Treichler, D., Wardlaw, A., Weiss, G. & Goeller, J. Experimental Investigation of Cavitation as a Possible Damage Mechanism in Blast-Induced Traumatic Brain Injury in Post-Mortem Human Subject Heads. J. Neurotrauma 34, 1589–1602, DOI: 10.1089/neu.2016.4600 (2017).

53. Laksari, K., Assari, S., Seibold, B., Sadeghipour, K. & Darvish, K. Computational simulation of the mechanical response of brain tissue under blast loading. Biomech. Model. Mechanobiol. 14, 459–472, DOI: 10.1007/s10237-014-0616-2 (2015).

54. Elster, N. et al. A critical review of experimental analyses performed on animals, post-mortem human subjects, and substitutes to explore primary blast-induced Traumatic Brain Injuries. Front. Mech. Eng. 9, 1185231, DOI: 10.3389/fmech.2023.1185231 (2023).

55. Clayton, E. H., Genin, G. M. & Bayly, P. V. Transmission, attenuation and reflection of shear waves in the human brain. J. The Royal Soc. Interface 9, 2899–2910, DOI: 10.1098/rsif.2012.0325 (2012).

56. Hosseini-Farid, M., Amiri-Tehrani-Zadeh, M., Ramzanpour, M., Ziejewski, M. & Karami, G. The Strain Rates in the Brain, Brainstem, Dura, and Skull under Dynamic Loadings. Math. Comput. Appl. 25, 21, DOI: 10.3390/mca25020021 (2020).

57. Garimella, H. T., Kraft, R. H. & Przekwas, A. J. Do blast induced skull flexures result in axonal deformation? PLOS ONE e0190881, DOI: 10.1371/journal.pone.0190881 (2018).

58. Miller, S. T. et al. Localizing Clinical Patterns of Blast Traumatic Brain Injury Through Computational Modeling and Simulation. Front. Neurol. 12, 547655, DOI: 10.3389/fneur.2021.547655 (2021).

59. Chafi, M. S., Ganpule, S., Gu, L. & Chandra, N. Dynamic response of brain subjected to blast loadings: Influence of frequency ranges. Int. J. Appl. Mech. 03, 803–823, DOI: 10.1142/S175882511100124X (2011).

60. Sahoo, D., Deck, C. & Willinger, R. Brain injury tolerance limit based on computation of axonal strain. Accid. Analysis & Prev. 92, 53–70, DOI: 10.1016/j.aap.2016.03.013 (2016).

61. Song, H. et al. Ultrastructural brain abnormalities and associated behavioral changes in mice after low-intensity blast exposure. Behav. Brain Res. 347, 148–157, DOI: 10.1016/j.bbr.2018.03.007 (2018).

62. Du, Z. et al. Mechanical mechanism and indicator of diffuse axonal injury under blast-type acceleration. J. Biomech. 156, 111674, DOI: 10.1016/j.jbiomech.2023.111674 (2023).

63. Sutar, S. & Ganpule, S. Investigation of wave propagation through head layers with focus on understanding blast wave transmission. Biomech. Model. Mechanobiol. 19, 875–892, DOI: 10.1007/s10237-019-01256-9 (2020).

64. Gupta, R. K., Tan, X. G., Somayaji, M. R. & Przekwas, A. J. Multiscale Modelling of Blast-Induced TBI Mechanobiology - From Body to Neuron to Molecule. Def. Life Sci. J. 2, 3, DOI: 10.14429/dlsj.2.10369 (2017).

65. Eslaminejad, A., Hosseini Farid, M., Ziejewski, M. & Karami, G. Brain Tissue Constitutive Material Models and the Finite Element Analysis of Blast-Induced Traumatic Brain Injury. Sci. Iran. 0, 0–0, DOI: 10.24200/sci.2018.20888 (2018).

66. Townsend, M. T., Alay, E., Skotak, M. & Chandra, N. Effect of Tissue Material Properties in Blast Loading: Coupled Experimentation and Finite Element Simulation. Annals Biomed. Eng. 47, 2019–2032, DOI: 10.1007/s10439-018-02178-w (2019).

