## Supplementary Information for "Biomechanical response of the human brain to low-intensity blast: a finite element study of single and repeated exposures"

**Supplementary Table S1.** Material parameters used in the APL Head-Neck model.

| Structure | Material model | LS-DYNA ID | $\rho_0$ (kg · m$^{-3}$) | Constitutive parameters |
|---|---|---|---|---|
| Anulus fibrosus | Visco-soft-tissue | `MAT_091` | 1,830 | $C_1 = 9.82\times10^{-4}$ GPa; $XK = 0.001099$ GPa; $XLAM = 1.0$; $AOPT = 2$; $A_3 = 1000.0$; $D_1 = 1000.0$; $MACF = 1$; $S_1 = 0.744$; $S_2 = 0.1098$; $S_3 = 0.0356$; $S_4 = 0.0251$; $S_5 = 0.0855$; $T_1 = 1.0$ ms; $T_2 = 10.0$ ms; $T_3 = 100.0$ ms; $T_4 = 1000.0$ ms; $T_5 = 1000000.0$ ms |
| Brain | Bergström–Boyce rubber | `MAT_269` | 1,000 | $K = 2.0$ GPa; $G = 5.494\times10^{-7}$ GPa; $G_v = 5.0\times10^{-6}$ GPa; $N = 92.353$; $N_v = 3.0$; $C = -0.234$; $M = 1.003$; $\dot{\gamma}_0 = 0.001$ ms$^{-1}$; $\hat{\tau} = 2.46\times10^{-6}$ GPa |
| Bridging veins | Elastic | `MAT_001` | 1,100 | $E = 0.03$ GPa; $\nu = 0.48$ |
| Cerebrospinal fluid | Viscoelastic | `MAT_006` | 1,000 | $K = 2.19$ GPa; $G_0 = 6.0\times10^{-7}$ GPa; $G_1 = 1.2\times10^{-7}$ GPa; $\beta = 0.08$ ms$^{-1}$ |
| Cortical bone | Elastic | `MAT_001` | 1,600 | $E = 10.0$ GPa; $\nu = 0.30$ |
| Dura mater | Elastic | `MAT_001` | 1,160 | $E = 0.0315$ GPa; $\nu = 0.35$ |
| Falx–tentorium | Elastic–plastic–thermal | `MAT_004` | 1,133 | $E_i = 0.036$ GPa; $\nu_i = 0.20$; $\alpha_i = 0.001$ K$^{-1}$; $\sigma_i = 1.0$ GPa |
| Flesh | Ogden rubber | `MAT_077_O` | 1,100 | $\nu = 0.498$; $\mu = 0.00148$ GPa; $\alpha = 8.1$ |
| Nucleus pulposus | Mooney–Rivlin rubber | `MAT_027` | 1,200 | $\nu = 0.49$; $A = 1.428\times10^{-4}$ GPa; $B = 3.57\times10^{-5}$ GPa |
| Sinuses | Elastic | `MAT_001` | 1,100 | $E = 0.0315$ GPa; $\nu = 0.35$ |
| Skin | Elastic | `MAT_001` | 1,130 | $E = 0.0167$ GPa; $\nu = 0.42$ |
| Trabecular bone | Elastic | `MAT_001` | 1,600 | $E = 0.6$ GPa; $\nu = 0.30$ |
| Vertebral body | Rigid | `MAT_020` | 2,000 | $E = 1.0$ GPa; $\nu = 0.30$ |
